# Building artificial plant cell wall on lipid bilayer by assembling polysaccharides and engineered proteins

**DOI:** 10.1101/2022.07.25.501355

**Authors:** Simona Notova, Nathan Cannac, Luca Rabagliati, Maeva Touzard, Josselin Mante, Yotam Navon, Liliane Coche-Guérente, Olivier Lerouxel, Laurent Heux, Anne Imberty

## Abstract

The cell wall constitutes a fundamental structural component of plant cells, providing them with mechanical resistance and flexibility. Mimicking that wall is a critical step in the conception of an experimental model of the plant cell. The assembly of cellulose/hemicellulose in the form of cellulose nanocrystals and xyloglucans as a representative model of the plant cell wall has already been mastered, however, those models lacked the pectin component. In this work, we used an engineered chimeric protein designed for bridging pectin to the cellulose/hemicellulose network, therefore achieving the assembly of complete cell wall mimics. We first engineered proteins, i.e. carbohydrate-binding module from *Ruminococcus flavefaciens* able to bind oligogalactorunan, resulting in high-affinity polygalacturonan receptors with Kd in the micromolar range. A Janus protein, with cell wall gluing property, was then designed by assembling this CBM with a *Ralstonia solanacearum* lectin specific for fucosylated xyloglucans. The resulting supramolecular architecture is able to bind fucose-containing xyloglucans and homogalacturonan ensuring high affinity for both. A two-dimension assembly of an artificial plant cell wall was then built first on synthetic polymer and then on supported lipid bilayer. Such artificial cell wall can serve as a basis for the development of plant cell mechanical models and thus deepen the understanding of the principles underlying various aspects of plant cells and tissues.

**Graphical abstract:** 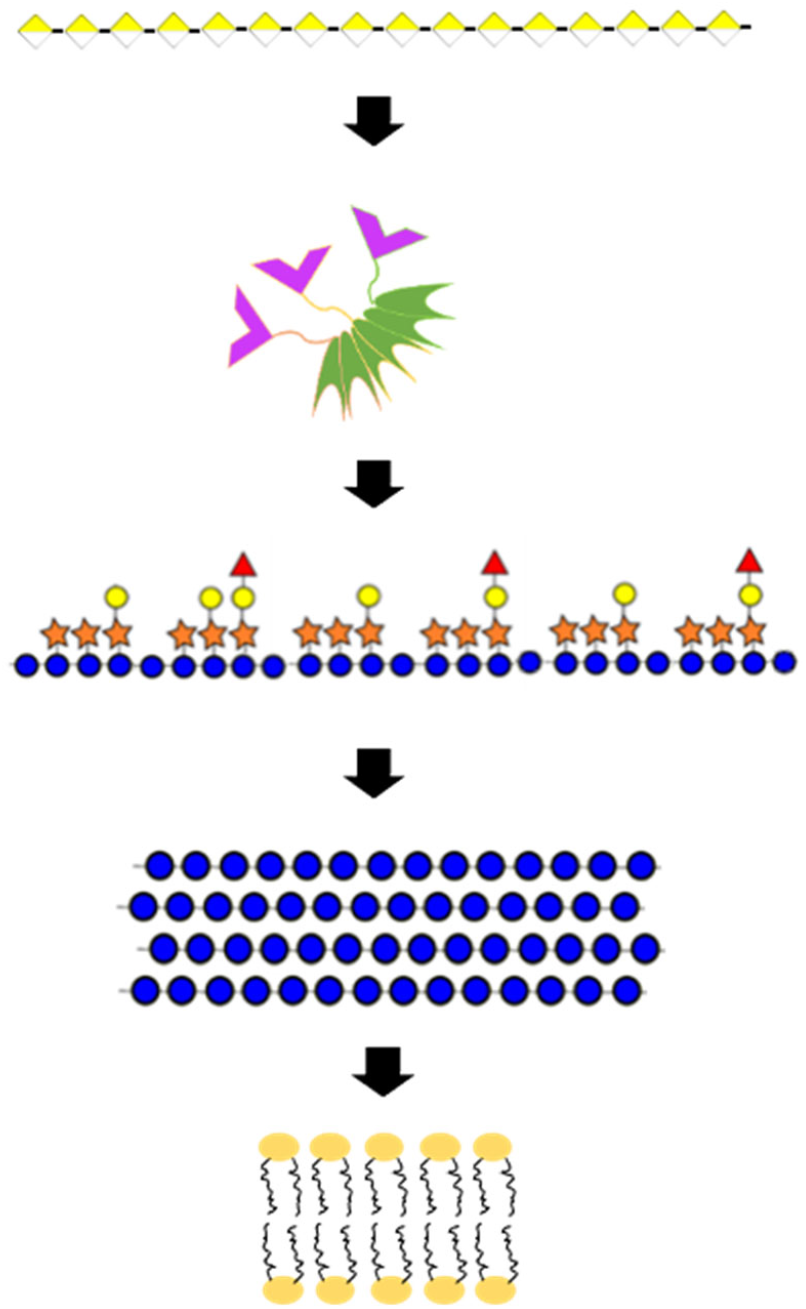

## Introduction

The composite material nature of the primary plant cell walls brings them their essential functions in controlling cell shapes, providing structural rigidity but also flexibility and establishing communications, exchange and also protecting barriers against pathogens (McNeil et al., 1984; B. Zhang et al., 2021). The primary cell walls consist mainly of cellulose microfibrils embedded in two other polysaccharides networks, hemicellulose, and pectin, with a low amount of proteins. Cellulose occurs as semi-crystalline microfibrils and forms a mesh-like structure around the cells. Hemicellulose, such as xyloglucan, adsorbs onto cellulose, crosslinks the microfibrils, and forms an entangled network that brings further strength and extensibility (Pauly et al., 1999). Pectins are about one-third of plant primary cell wall polysaccharides and the major charged polysaccharides of this complex system. They are mainly composed of homogalacturonans which form a network in close contact but independent of cellulose and through phase separation processes, are a driving force for cell wall expansion and morphogenesis (Haas et al., 2020). Pectin’s homogalacturonan nanofilament expansion drives morphogenesis in plant epidermal cells (Haas et al., 2021). Both networks gave a complex mechanical behavior to the plant cell wall essential to provide elasticity, plasticity, and deformability upon mechanical stress (Y. Zhang et al., 2021). Although present in lower amounts, cell wall proteins have essential roles, and are prone to be involved in cell wall dynamics like expansins, or having structural functions that cross-link the component of the cell wall and should participate in its mechanical properties like extensins (Jamet et al., 2006).

The remarkable composition of plant cell walls inspired the design of biomimetic materials, that are environmentally friendly and have unique properties in strengths, lightness, and resistance (Teeri et al., 2007). Applications range from pharmaceutics with polysaccharide-based vesicles for encapsulating active compounds to the coating, or even packaging. Recently, the engineering of multilayer microcapsules based on polysaccharides nanomaterials resulted in vesicles with various applications (Lombardo & Villares, 2020). For the formation of such vesicles, cellulose nanomaterials are often the main component in a bio-mimetic approach, with the addition of xyloglucan or pectin. The most recent study succeeded in assembling cellulose microfibrils and pectins on lipid-based vesicles and therefore reached one step closer to the construction of artificial plant cells (Paulraj et al., 2020). Even though the individual components of the plant cell wall found their applications, the construction of synthetic plant cell walls remains a challenge.

Biomimetic cell walls can also be obtained in the form of film, i.e., supported multilayer assembly that can be analyzed by a variety of techniques. Such films were first obtained based on cellulose nanocrystals (CNCs) that can be easily obtained from plant material with strong acid treatment, hydrolyzing the amorphous regions of cellulose and releasing intact crystalline blocks (Habibi et al., 2010; Rånby, 1949). CNCs have high colloidal stability and represent an excellent model for mimicking the naturally occurring cellulose microfibrils. Multilayers of CNCs and xyloglucan (hemicellulose) can be obtained relatively easily due to the strong interactions, mainly driven by hydrophobic interaction occurring between the nonpolar faces of the glucose chain present in both polysaccharides (Hanus & Mazeau, 2006; Hayashi et al., 1994; Lopez et al., 2010; Mazeau & Charlier, 2012). In order to mimic plant cell wall, a lipidic component is necessary. Supported lipid bilayer (SLB) is a widely used and characterized model of cell membranes (McConnell et al., 1986; Richter & Brisson, 2005). Interaction between CNCs and SLB is the crucial step for an artificial plant cell wall and has also been described after tuning of parameters and the resulting multilayer has been fully characterized (Y. Navon et al., 2020). Additionally, the addition of xyloglucan to this lipid-supported nanocellulose film has also been reported (Yotam Navon, 2020).

The next step is therefore to include a layer of pectin on the multilayer. Pectins are a family of complex, charged and functionalized polysaccharides, the simplest model available is homogalacturonan (HG) i.e., a linear polymer of α1-4 linked galacturonic acid (GalA). Unmethylated HG can be obtained from different plant sources after enzymatic treatment. The adsorption of such soluble and charged polysaccharides on CNCs-xyloglucan film is a difficult process, partly due to the competition between xyloglucan and pectin for the binding to cellulose (Zykwinska et al., 2008). In the cell wall, a tiny amount of proteins is present, and extensins are known to link the pectin (Sede et al., 2018). We, therefore, propose to engineer protein glue that would stabilize the interaction between hemicellulose and pectin for the assembly of an artificial cell wall.

Proteins able to bind to oligo or polysaccharides without modifying them belong to two main families. Lectins are generally multivalent proteins with several carbohydrate-binding sites and exist with a variety of sequence and fold (Bonnardel et al., 2019). On the other hand, carbohydrate-binding modules (CBMs) are monovalent protein domain (Lombard et al., 2014), generally associated with a carbohydrate-active enzyme. Many of them play a role in biomass degradation, and are specific for plant polysaccharides (Gilbert et al., 2013). While CBMs have low affinity for glycans, it has been demonstrating that associating two of them by flexible linkers results in 2 to 3 order of magnitude enhancement of the affinity (Connaris et al., 2009; J. Ribeiro et al., 2016). Such synthetic biology approach was also used for building a bi-specific Janus lectin, assembling six fucose binding domains with three sialic acid binding domain (J. Ribeiro et al., 2016). This construct was able to crosslink vesicles decorated with the appropriate glycolipids, but also to assemble a multilayer of glycosylated multivalent compounds, forming a multilayer assembly.

We propose here to design a chimeric lectin with one face able to bind hemicellulose and one face specific for homogalacturonan. Such construct should be able to glue the xyloglucan/cellulose cement deposited on supported lipid, with the pectin matrix added above, therefore providing a more complete model of the whole plant cell wall.

## Results

### Selection of biomolecules for the building of artificial cell wall

The construction of artificial cell walls involved the polysaccharides displayed in Figure 1, i.e. cellulose nanocrystals (CNC), fucosylated xyloglucan (FXG) as representative hemicellulose polysaccharide and linear homogalacturonan (HG) consisting of galacturonic acid (GalA) linked by α1-4 linkages which represents the simplest model for pectin. The preparation, purification and characterization of these three polysaccharides components are described in the material and methods section.

**Figure 1:**
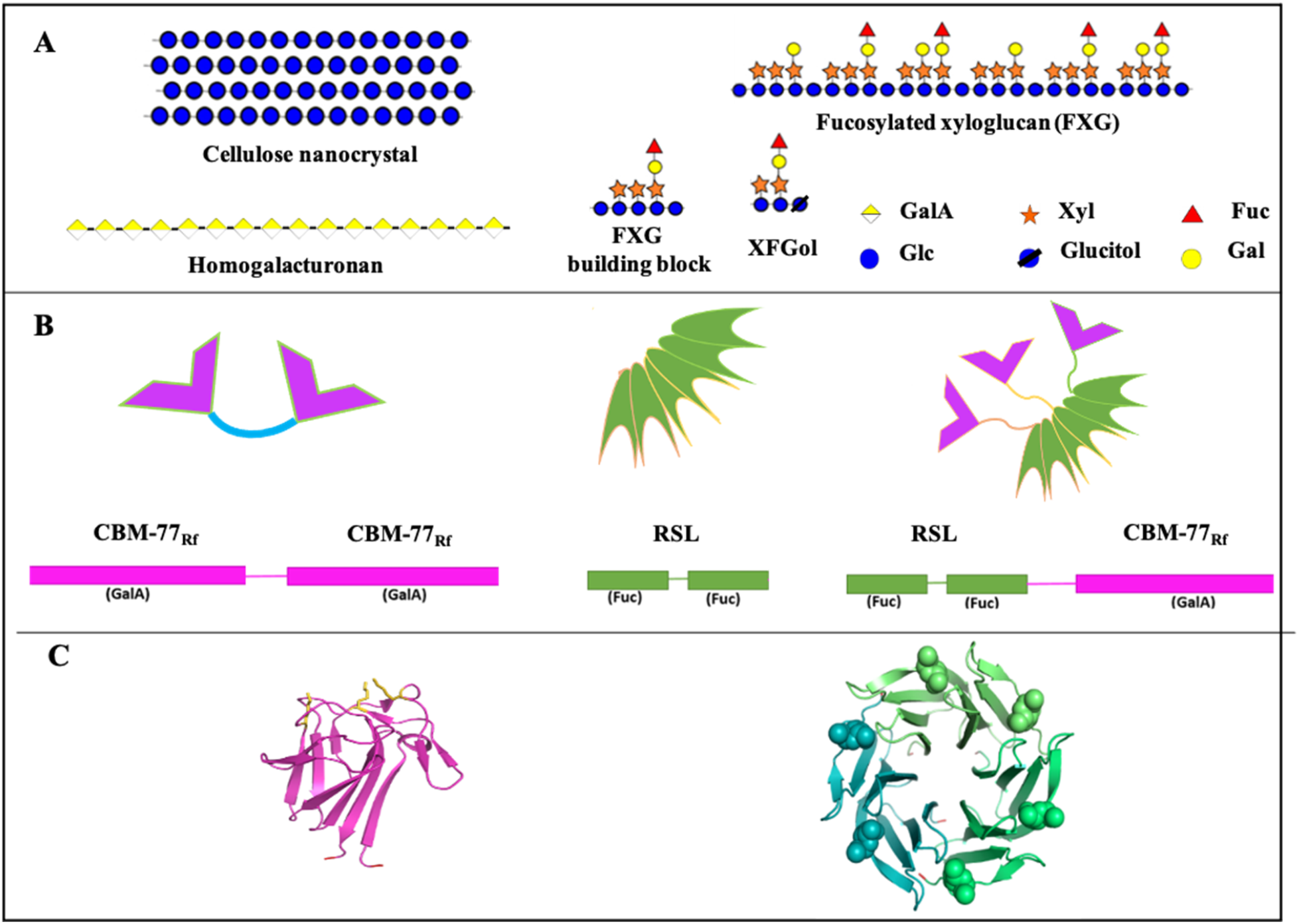
Biomolecules used in the building of artificial plant cell walls. A: Polysaccharides such as crystalline nanocellulose (CNC), homogalacturonan (HG) from pectin, and fucosylated xyloglucan (FXG) from hemicellulose, together with a schematic representation of monosaccharides in agreement with SNFG nomenclature (Neelamegham et al., 2019). B: Schematic representation and peptide organization of engineered diCBM77_Rf,_ RSL and Janus RSL-CBM77_Rf_. C: Three-dimensional structure of GalA-specific monomeric CBM77_Rf_ from *Ruminococcus flavefaciens* (PDB 5FU5) and fucose-specific trimeric RSL from *Ralstonia solanacearum* (PDB 2BT9) with protein represented by ribbon and fucose represented by spheres.

In order to create an interface able to crosslink hemicellulose and pectin, we aimed at building a chimeric lectin that binds both polysaccharides. In the first step, we searched for carbohydrate-binding modules (CBMs) with specificity for HG in databases such as CAZY (Drula et al., 2022) and CBM-Carb (https://cbmdb.glycopedia.eu/). This resulted in the identification of two previously characterized proteins, YeCBM32 from *Yersinia enterolitica* (Abbott et al., 2007) and CBM77_RfPL1/9,_ here referred as CBM77_Rf,_ from *Ruminococcus flavefaciens* (Venditto et al., 2016). Both of them are modules associated with pectin lyases involved in the degradation of plant cell walls and display a strong affinity for HG and oligogalacturonans with a size greater than six GalA residues. After testing for possibility of recombinant production, CBM77_Rf_ was selected for the project, first for engineering it as a dimer in order to check if sufficient avidity can be obtained for pectin, and then as a multivalent chimeric construct (Figure 1). In order to create a chimeric lectin with two specificities, a hemicellulose specific-binding protein has to be associated with CBM77_Rf_. It was previously demonstrated that the fucose binding lectin from bacteria *Ralstonia solanacearum*, presents strong affinity for oligomers of fucosylated xyloglucans (Kostlanová et al., 2005). This lectin associates as a trimer and possesses six fucose binding sites due to the presence of tandem repeats in each monomer. RSL and CBM77_Rf_ were then associated as chimeric lectin through addition of a linker peptide.

### Engineered lectins with specificity for plant cell wall polysaccharides

#### Design and production of divalent CBM77_Rf_ with high avidity for homogalacturonan

The engineering of CBMs was already proven as an efficient tool for increasing protein valency and affinity (Connaris et al., 2009; J. Ribeiro et al., 2016). Using similar strategy, a dimeric CBM77_Rf_ (diCBM77_Rf_) was designed in order to confirm that multivalent organization has the potential to increase the affinity for HG. The protein was engineered as a tandem repeat of two CBM77_Rf_ connected through flexible linker ALNGSELGSGSGLSSLGEYKDI which was previously used in the design of another divalent CBM, diCBM40 (J. Ribeiro et al., 2016). diCBM77_Rf_ is composed of 270 amino acids and was fused with N-terminal His-tag associated with TEV (tobacco etch virus) protease site. The protein was recombinantly produced in soluble form in bacteria *E. coli* BL21(DE3) and purified by immobilized metal ion chromatography followed by His-tag cleavage by TEV protease. After the TEV cleavage, the protein consisted of 251 AA with an estimated molecular weight of 25.7 kDa (Figure 2A).

**Figure 2:**
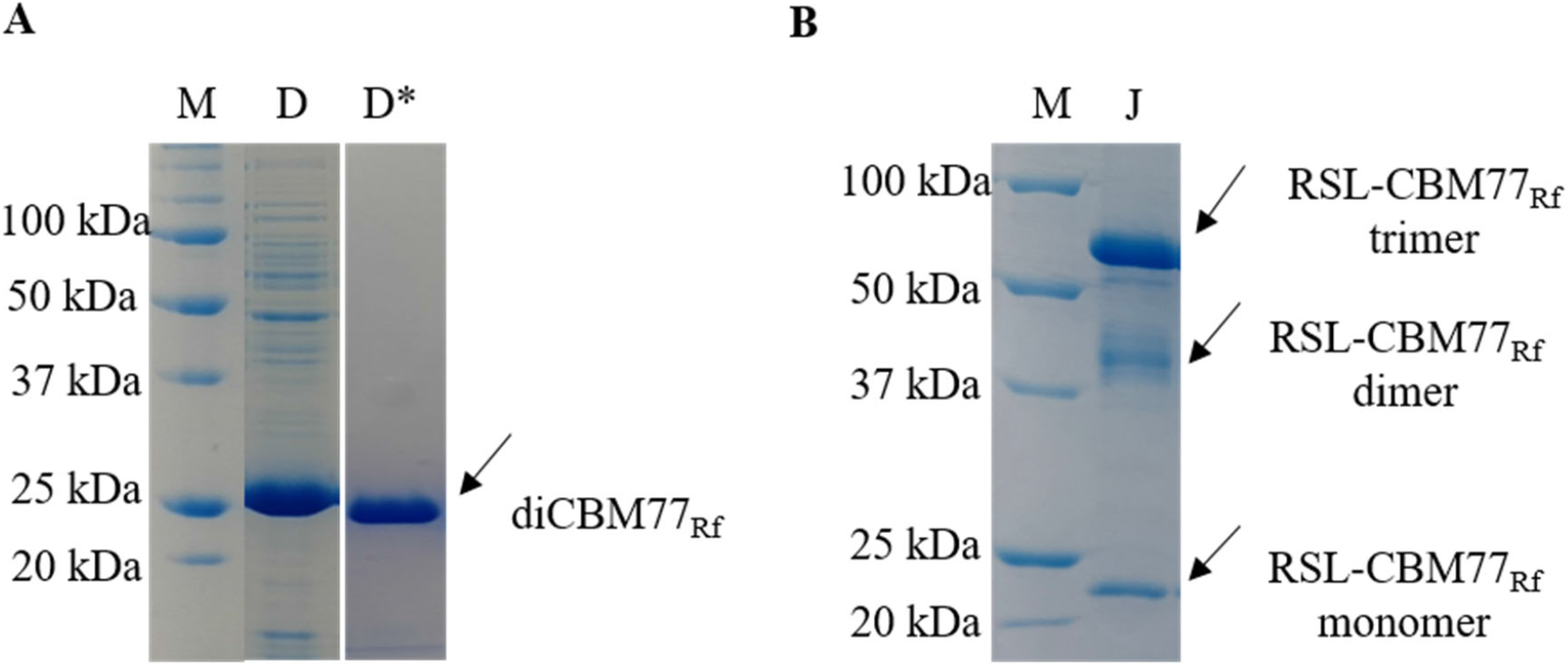
SDS page analysis of diCBM77_Rf_ and Janus lectin RSL-CBM77_Rf_. The protein samples were analyzed under denaturing conditions on 12% polyacrylamide gel. M-protein marker, D-diCBM77_Rf_, D*-diCBM77_Rf_ cleaved by TEV protease, J-Janus lectin RSL-CBM77_Rf_. A) monomeric diCBM77_Rf_ with an estimated size of 27.9 kDa. The impurities were eliminated once the protein was cleaved with TEV protease and the His-tag was removed, B) Janus lectin RSL-CBM77_RF_ appears to be trimeric even in denaturation conditions, the trimerization is a result of RSL stable oligomerization. The molecular weight was estimated as 66 kDa, 44 kDa, and 22 kDa corresponding to trimeric, dimeric, and monomeric RSL-CBM77_Rf_, respectively.

#### Design and production of a Janus lectin with the ability to cross-link pectin and xyloglucan

The strategy to obtain a Janus lectin was to connect the gene coding for RSL monomer (N-terminus) with the one of CMB-77 domain (C-terminus) through additional oligonucleotide sequence coding for flexible linker (GGGGSGGGGS) as previously used for engineering of Janus lectin (J. P. Ribeiro et al., 2018). The resulting protein consists of 214 amino acids and was recombinantly produced in soluble form in *Escherichia coli* KRX and subsequently purified by affinity as previously described for lectin RSL (Kostlanová et al., 2005). RSL-CBM77_Rf_ has an estimated molecular weight of 22.1 kDa however, SDS-PAGE analysis showed mostly the presence of protein with the estimated size of 66 kDa suggesting that RSL-CBM77_Rf_ assembles as a stable trimer that does not separate even during denaturation (Figure 2B). The oligomerization is under the control of RSL and the protein displays three HG binding sites site on one face and six fucose binding sites on the other face.

### Binding to plant cell wall oligo- and polysaccharides

#### diCBM77_Rf_ binding to oligo- and polygalacturonate

The efficiency of diCBM77_Rf_ for binding to pectin fragments was assayed by ITC, with the comparison of affinity towards oligomers and polysaccharides. When titrating diCBM77_Rf_ with hepta-galacturonic acid (GalA_DP7) large exothermic peaks were obtained, indicating exothermic binding (Figure 3A). The affinity was in the millimolar range, and the sigmoid curve could not be obtained. On the opposite, binding to homogalacturonan (HG), resulted in strong avidity, with an affinity value of 780 nM (Figure 3B). The stoichiometry (N) indicated the binding of 14 diCBM77_Rf_ units per HG polysaccharide. Since the polysaccharide sample was characterized with a molecular weight (MW) of approx. 35 kDa, corresponding to a degree of polymerization (DP) of 180 GalA, each diCBM77_Rf_ occupies 14 GalA residues (either linear or cross-link), i.e., seven GalA per CBM77_Rf_ monomer, therefore GalA_DP7 can be considered as an effective binding unit. When calculating affinity and thermodynamic contribution per CBM77_Rf_ (i.e., per heptamer of GalA), the affinity is 20 µM. The engineering of a diCBM77_Rf_, therefore, resulted in multiplying the avidity by 60-fold when compared to monomeric diCBM77_Rf_ for the equivalent substrate.

**Figure 3:**
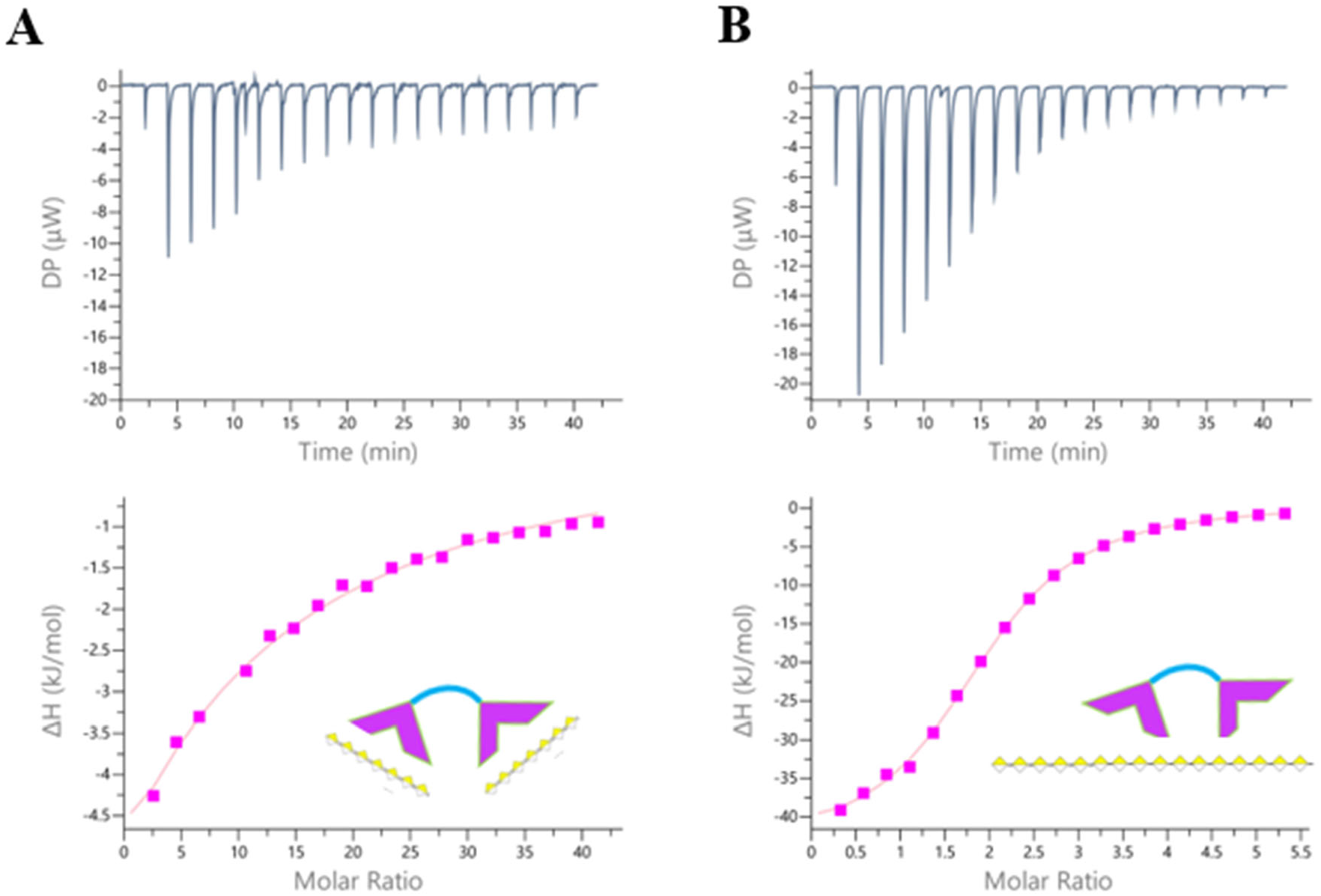
ITC data for the binding of diCBM77_Rf_ on pectin oligosaccharide Gal_DP7 (A) and polysaccharide HG (B). The figures were prepared using ligand concentration per effective binding unit, i.e., GalA_DP7. Top: thermogram obtained by injecting oligo or polysaccharides into the protein solution. Bottom: integration of peaks (rectangles) and obtained fit (line). ITC conditions: A) 50 μM diCBM77_Rf_ in buffer A with 10 mM GalA_DP7 in 20 mM Tris pH 7.5, B) 100 μM diCBM77_Rf_ in buffer A with 100 μM HG in buffer A.

#### Janus lectin RSL-CBM77_Rf_ binding to plant cell wall oligo and polysaccharides

The Janus lectin, consisting of a trimer of RSL-CBM77_Rf_, was tested by ITC for its binding capacity toward polysaccharides and oligosaccharides. RSL-CBM77_Rf_ binds to GalA_DP7, indicating that the addition of RSL peptide at the N-terminal extremity of CBM77_Rf_ does not alter its activity. In fact, trimerization enhanced the affinity of CBM77_Rf_ towards GalA_DP7, with a Kd of 103 µM, therefore 10 times higher than what was observed for diCBM77_Rf_ (Figure 4A). Similarly binding to HG is more efficient for Janus lectin RSL-CBM77_Rf_ than for diCBM77_Rf_, resulting in a very strong affinity of 211 nM (Figure 4B). When considering GalA_DP7 as the effective binding unit for CBM77_Rf_, which again corresponds closely to the observed stoichiometry (approx. 10 Janus lectin trimer per HG polysaccharides), an affinity of 4.8 µM is obtained, approximately 20 times better than for the oligomer in solution. The affinity for HG is four time better for the Janus lectin than for the diCBM77_Rf_. Analysis of the thermodynamic contribution indicates that the gain in affinity is related to a lower entropy barrier. It may be that the architecture of Janus lectin brings the three CBMs close to each other, resulting in a much more efficient binding of the substrate, and therefore stronger affinity as described for other multivalent systems (Dam & Brewer, 2002). However, decrease in entropy barrier is also observed when comparing Janus lectin RSL-CBM77_Rf_ and diCBM77_Rf_ binding to the oligosaccharide, indicating that other effects also play a role.

**Figure 4:**
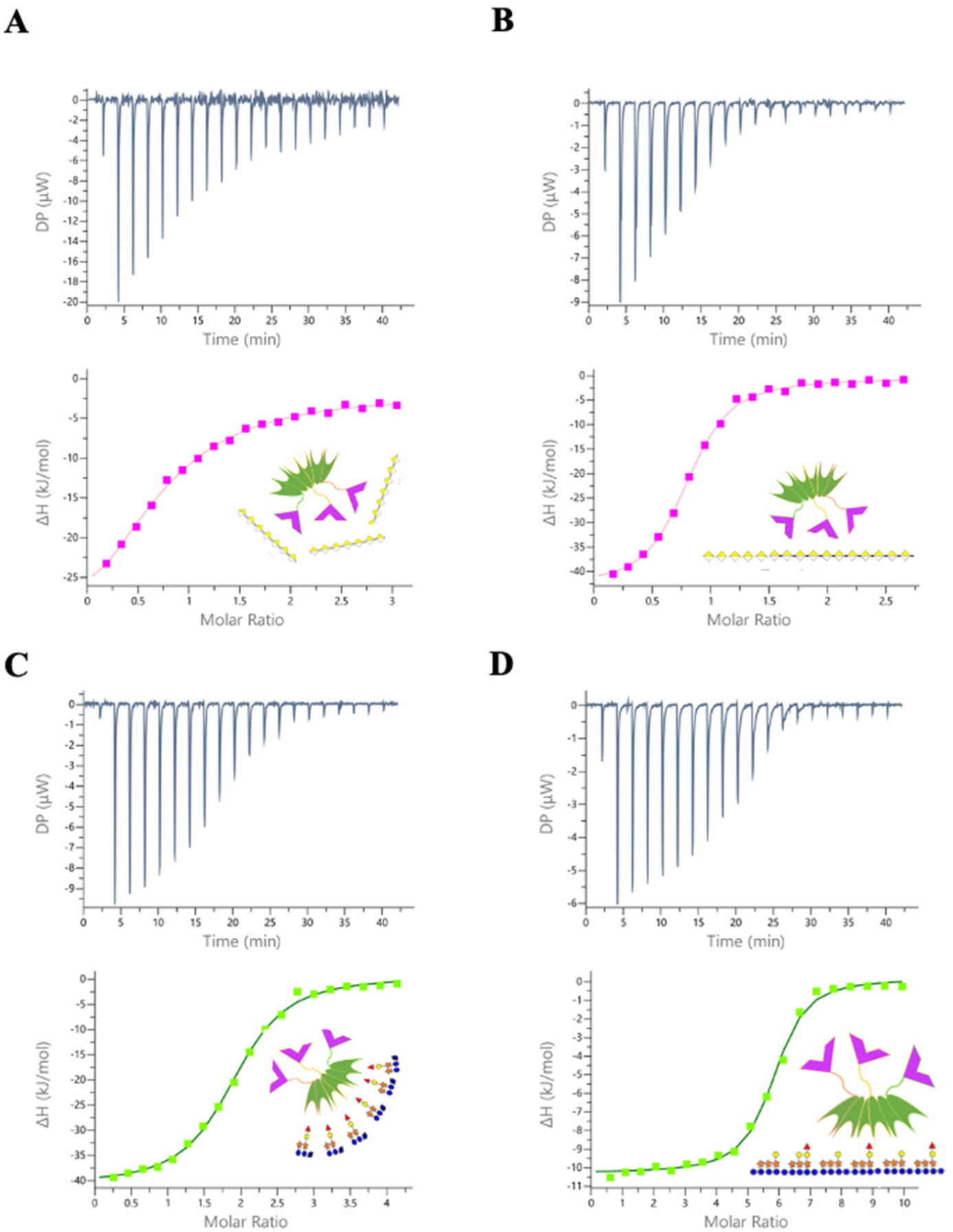
Comparison of ITC isotherms of RSL-CBM77_Rf_ obtained by titration with oligosaccharides and polysaccharides. Top: thermograms are obtained by injecting oligo or polysaccharides into the solution of protein. The figures were prepared using ligand concentration per effective binding unit, i.e., seven GalA stretch motif for HG or ten monosaccharide building block for apple FXG. Bottom: integration of peaks (rectangles) and obtained fit (line). ITC conditions: A) 204 µM RSL-CBM77_Rf_ in buffer A with 3 mM GalA_DP7 oligosaccharide in 20 mM Tris pH 7.5, B) 100 µM RSL-CBM77_Rf_ in buffer A with 100 µM HG in buffer A, C) 50 µM RSL-CBM77_Rf_ in buffer A with 1 mM XFGol oligosaccharide in buffer A, D) 50 µM RSL-CBM77_Rf_ in buffer A with 75 µM apple FXG in buffer A.

As for binding to fucosylated oligosaccharides, each Janus lectin RSL-CBM77_Rf_ binds with strong affinity to mono-fucosylated octasaccharide XFGol (Kd = 3.9 µM) (Figure 4C). The observed stoichiometry of 1.88 is in agreement with the occurrence of two fucose binding site per protein monomer (i.e. six per Janus lectin trimer). The obtained affinity is very similar to that measured for RSL for control experiments (Figure 5) and also corresponds to previously published results with oligosaccharide XXFG with the affinity of 2.8 µM obtained by surface plasmon resonance (Kostlanová et al., 2005). The Janus lectin displays much stronger avidity for apple fucosylated xyloglucan (FXG) than for oligosaccharides, with a measured Kd of 51 nM (Figure 4D). The measured stoichiometry of 0.168 corresponds to six RSL monomers (each with two fucose binding sites) per polysaccharide FXG, indicating the presence of a minimum of 6 to 12 fucose residues per polymer. This is in agreement with the known structures of apple pomace xyloglucan (Watt et al., 1999) and with the evaluation by gel permeation chromatography (GPC) and decomposition analysis (see material and methods). GPC revealed the estimated molecular weight of apple FXG of 55 kDa which corresponds to approximately 300 monosaccharide residues and decomposition analysis confirmed that 5% of this polymer is fucose, which corresponds to 15 monosaccharide units (Table 4). The affinity of the lectin was therefore recalculated per fucose binding unit with the value of 620 nM. The stoichiometry 2 indicates that both binding sites of the RSL part of monomeric RSL-CBM77_Rf_ were occupied with the fucose. The affinity was also determined per building block (i.e. xyloglucan repeat of XFG or XXG - see Fig1A). Their average composition of ten monosaccharide units would result in 30 such building blocks per FXG polysaccharide. The observed stoichiometry of 5.4 building block for each monomeric RSL-CBM77_Rf_ confirmed that fucose is present in every 2 or 3 building blocks of the polymer FXG.

**Figure 5:**
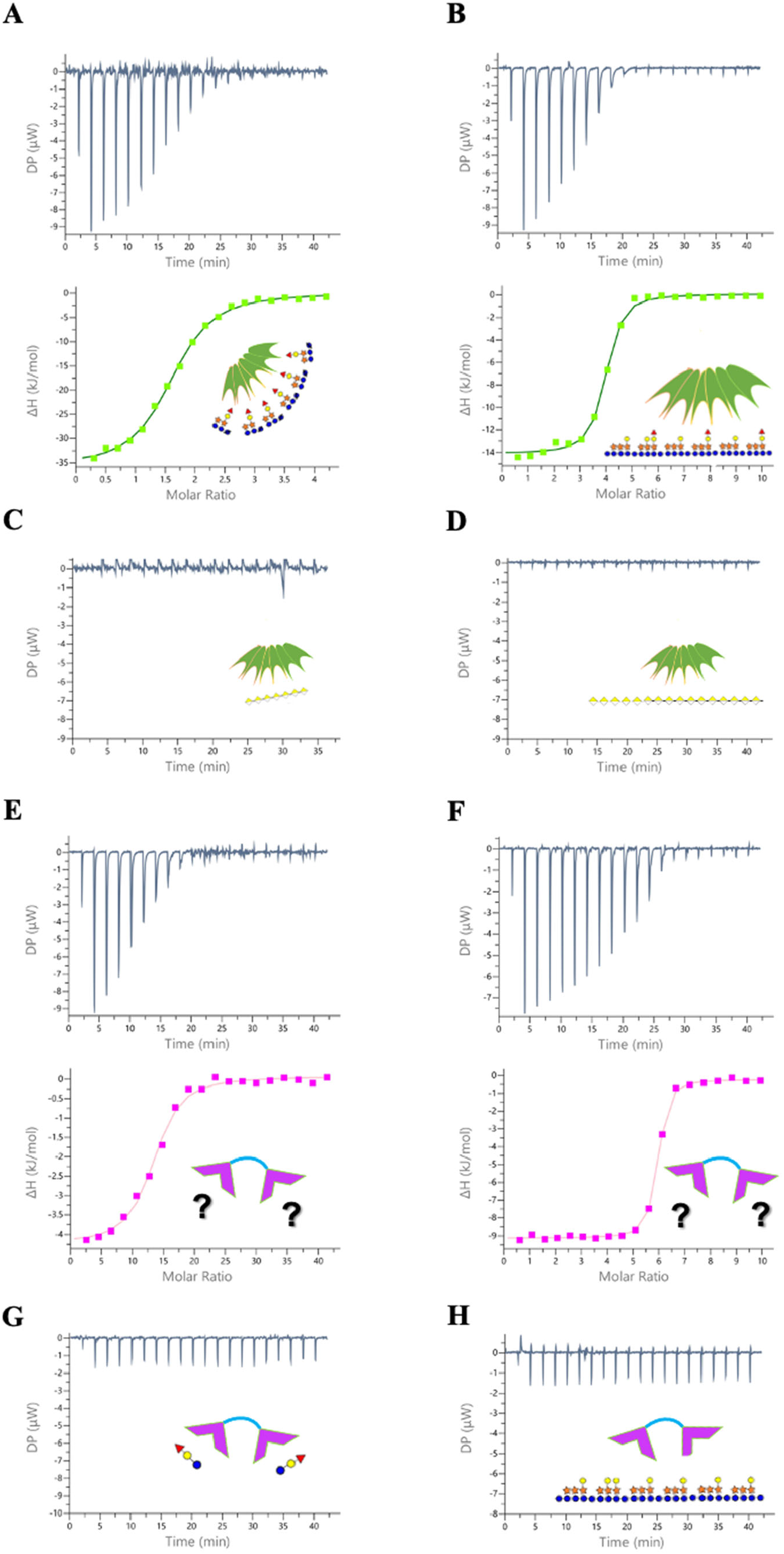
Control ITC experiments. ITC conditions: A) 50 µM RSL in buffer A with 1 mM XFGol in buffer A, B) 50 µM RSL in buffer A with 100 µM apple FXG in buffer A, C) 50 µM RSL in buffer A with 10 mM GalA_DP7 oligosaccharide in 20 mM Tris pH 7.5, D) 50 µM RSL in buffer A with 50 µM HG in buffer A, E) 50 µM diCBM77_Rf_ in buffer A with 10 mM XFGol in buffer A, F) 50 µM diCBM77_Rf_ in buffer A with 50 µM apple FXG in buffer A, G) 50 µM diCBM77_Rf_ in buffer A with 10 mM 2-fucosyl lactose in buffer A, H) 50 µM diCBM77_Rf_ in buffer A with 4 µM tamarin nonfucosylated XG in buffer A. Top: thermograms obtained by injecting of oligo or polysaccharides into the protein solution. Bottom (except C, D): integration of peaks (rectangles) and obtained fit (line).

#### Control experiments indicating presence of polysaccharide contaminant

Control experiments were performed with both RSL and diCBM77_Rf_ in order to check the specificity.

RSL alone binds as expected on XFGol and FXG polymer (Figure 5A, B). The binding to the oligomers was identical than for Janus lectin. For binding to FXG polymer we determined the affinity and stoichiometry per fucose unit and FXG building block. As shown in Table 2 in both cases, the affinity of RSL and RSL-CBM77_Rf_ are comparable. The difference is however in stoichiometry per FXG building block, suggesting the binding of 4 building blocks per RSL monomer. Nevertheless, these finding just confirmed our hypothesis and calculation about the presence of fucose 50% of building block repeats of FXG polymer.

**Table 1:** Thermodynamic data expressed as a function of the carbohydrate ligand concentration, either as the whole molecule or as the “effective binding unit” concentration of the ligand, i.e., seven GalA stretch motif for HG, one fucosylated motif or ten monosaccharide building block for apple FXG. N indicates the measured ligand/protein stoichiometry.

| Prot. | Ligand | N | K<sub>d</sub><br>(μM) | ΔG<br>(kJ/mol) | ΔH<br>(kJ/mol) | -TΔS<br>(kJ/mol) |
| --- | --- | --- | --- | --- | --- | --- |
| **diCBM77<sub>Rf</sub>** |  |  |  |  |  |  |
| | GalA_DP7 | 2<sup>a</sup>(1)<sup>b</sup> | 1280 ± 100 | -16.5 | -63.7 ± 1.3 | 47.1 |
| | HG | 0.074 ± 0.00<br>(0.037)<sup>b</sup> | 0.78 ± 0.02 | -34.8 | -1135 ± 15 | 1110 |
| | HG<br>(per GalA_DP7) | 1.91 ± 0.01<br>(0.95)<sup>b</sup> | 20 ± 0.4 | -26.8 | -44.2 ± 0.45 | 17.3 |
| **Janus RSL-CBM77<sub>Rf</sub>** |  |  |  |  |  |  |
| | GalA_DP7 | 0.63 ± 0.01 | 103 ± 15 | -22.8 | -53 ± 4 | 30.2 |
| | HG | 0.031 ± 0.001 | 0.211 ± 0.01 | -38.1 | -1125 ± 5 | 1085 |
| | HG<br>(per GalA_DP7) | 0.78 ± 0.01 | 4.84 ± 0.31 | -30.0 | -43.7 ± 0.2 | 13.7 |
| | XFGol | 1.88 ± 0.01 | 3.9 ± 0.4 | -30.9 | -40.1 ± 0.1 | 9.2 |
| | FXG | 0.168 ± 0.01 | 0.051 ± 0.02 | -41.8 | -327.5 ± 7.5 | 285.3 |
| | FXG<br>(per Fuc unit) | 2.0 ± 0.1 | 0.62 ± 0.2 | -35.6 | -27.3 ± 0.6 | 8.3 |
| | FXG<br>(per building block) | 5.37 ± 0.26 | 1.65 ± 0.58 | -33.2 | -10.3 ± 0.3 | 22.9 |

<sup>a</sup>: value fixed during fitting procedure; <sup>b</sup>: Stoichiometry ligand/protein calculated per CBM77<sub>Rf</sub> monomer for comparison

**Table 2:** Thermodynamic data of control experiments. The data were prepared in the same way as previously shown in Table 1.

| Prot. | Ligand | N | Kd<br>(μM) | ΔG<br>(kJ/mol) | ΔH<br>(kJ/mol) | -TΔS<br>(kJ/mol) |
| --- | --- | --- | --- | --- | --- | --- |
| **diCBM77<sub>Rf</sub>** |  |  |  |  |  |  |
| | XFGol | 11.85 ± 0.95 | 21.6 ± 1.9 | -26.6 | -3.96 ± 0.37 | 22.7 |
| | FXG | 0.12 ± 0.00 | 0.013 ± 0.01 | -45.5 | -427.5 ± 0.5 | 381.5 |
| | FXG<br>(per building block) | 5.72 ± 0.02 | 0.62 ± 0.3 | -35.9 | -8.9 ± 0.0 | 27.1 |
|  | 2-fucosyl lactose |  | nb |  |  |  |
|  | XG |  | nb |  |  |  |
| **RSL** |  |  |  |  |  |  |
|  | GalA_DP7 |  | nb |  |  |  |
|  | HG |  | nb |  |  |  |
| | XFGol | 1.69 ± 0.1 | 5.1 ± 1.1 | -30.3 | -36.9 ± 0.9 | 6.6 |
| | FXG | 0.163 ± 0.01 | 0.46 ± 0.01 | -41.9 | -343 ± 3 | 301.5 |
| | FXG<br>(per Fuc units) | 2 ± 0.02 | 0.55 ± 0.08 | -35.7 | -28.6 ± 0.3 | 7.2 |
| | FXG<br>(per building block) | 3.9 ± 0.1 | 1.1 ± 0.2 | -34.0 | -14.3 ± 0.1 | 19.8 |

nb: no binding observed

In order to prove the specific mode of interaction for each polysaccharide control experiments were performed by ITC as well. RSL did not bind to neither oligosaccharide GalA_DP7 nor the polymer HG (Figure 5C, D). Similarly, diCBM77_Rf_ was tested with oligosaccharide XFGol and apple FXG. Surprisingly, a strong binding was observed with both solutions (Figure 5E, F). More investigations were conducted and diCBM77_Rf_ was tested with 2-fucosyllactose, corresponding to terminal disaccharide of XFGol, and non fucosylated xyloglucan from tamarin seeds. The experimental conditions were identical as those for XFGol and FXG and as shown on Fig. 5G and 5H, in this case, no binding was observed. These findings brought us to the conclusion that diCBM77_Rf_ does not interact with FXG since it does not bind to pure fucosylated oligosaccharide, nor to the nonfucosylated xyloglucan.

It is therefore very likely that the high affinity binding observed for diCBM77_Rf_ for XFGol and FXG polysaccharide is due the presence of a contaminant. Indeed, the stoichiometry of XFGol binding to diCBM77_Rf_ (n=12) indicates that only 8-16 % of the ligands are active. Oligosaccharide XFGol (Elicityl) was characterized with a purity of 85% (provided by the manufacturer) and therefore 15% of the sample composition is unknown. Similarly, apple FXG was characterized by decomposition analysis confirming the presence of additional monosaccharides such as arabinose, mannose, and rhamnose (Table 4). Since XFGol was produced from the same source as FXG (apple pomace), the same contamination might be present in both samples. The presence of pectic fragments in FXG polysaccharide is however not so surprising as the existence of covalent linkages between pectin and xyloglucan was first proposed in the plant cell wall model of Keegstra *et al*. (Keegstra et al., 1973) and further demonstrated by Thompson and Fry in suspension-cultured rose cells (Thompson & Fry, 2000). Further investigation is needed in order to determine the actual essence of the interaction. Nevertheless, this observation is not a barrier to the construction of artificial cell wall as the construction is performed sequentially.

### Building of supported synthetic plant cell wall

A multilayer model of plant cell walls was prepared and analyzed qualitatively through Quartz Crystal Microbalance with energy Dissipation monitoring technique (QCM-D). QCM-D measurements were used to monitor the adsorption of CNCs, polysaccharides, and proteins, on the surface of a silica-coated quartz crystal that was first functionalized with a thin film of polyethylene-imine (PEI), a branched cationic polymer which was previously demonstrated to provide a strong basis for adhesion of CNC (Y. Navon et al., 2020) or with supported lipid bilayer (SLB).

In the QCMD experiments, the adsorption or binding of particles, polymer or molecules will induce both a change in the frequency and dissipation. Concerning the frequency shift, it can be related to the change in mass by the Sauerbrey relationship (Reviakine et al., 2011) considering a rigid layer :

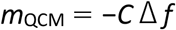

Regarding the changes in energy dissipation for each layer, the ratio (ΔD/-Δf) measuring the change in energy dissipation per surface-coupled mass unit characterizes the softness of the film, and it was demonstrated that a homogeneous film should be considered as rigid only for (Δ*D*/-Δ*f*) << 4 × 10^−7^ Hz^-1^. (Reviakine et al., 2011)). Above this limit, the films can be considered as soft and highly hydrated films.

Figure 6A displays the evolution of resonance frequency (related to the change in mass) and energy dissipation of the shear oscillation (related to the viscoelastic properties of the oscillating mass) for each layer deposited on the quartz crystal covered with PEI. Quantitative results are reported in Table 3.

**Figure 6:**
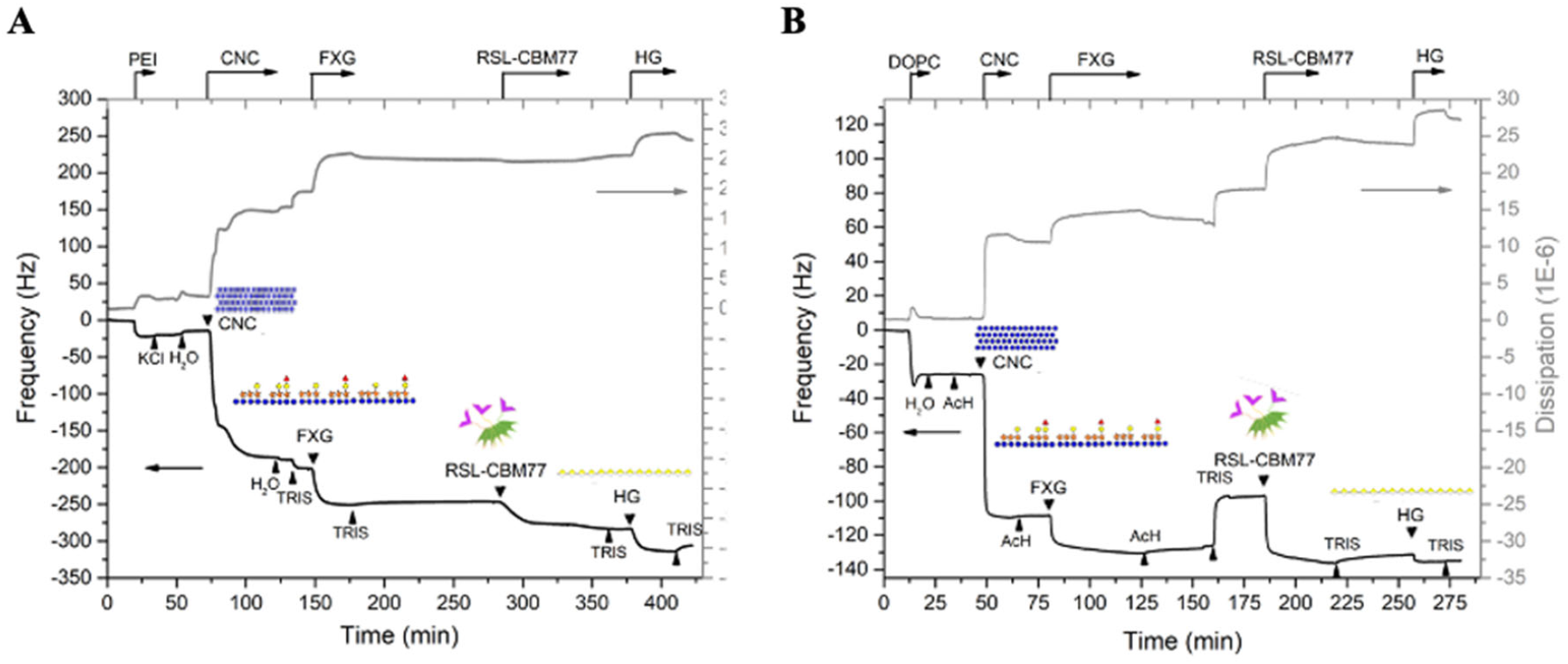
Frequency and dissipation variation over time of the quartz surface for the 7^th^ overtone with layer-by-layer stacking. **A:** PEI (1g.L^-1^) in 500mM KCl, CNC (1g.L^-1^) and FXG (1g.L^-1^) in pure water. This was followed by rinsing with 20mM Tris (pH=7.5), 100mM NaCl, an injection of RSL-CBM77_RfPL1/9_ (0.2, 2, and 10µM), and injection of HG in the same buffer. **B:** DOPC SLB (0.1g.L^-1^) in water, CNC (1g.L^-1^) and FXG (1g.L^-1^) in 100mM acetate. This was followed by rinsing with 20mM Tris (pH=7.5), 100mM NaCl, injection of RSL-CBM77_Rf_, and purified HG in the same buffer

**Table 3:**
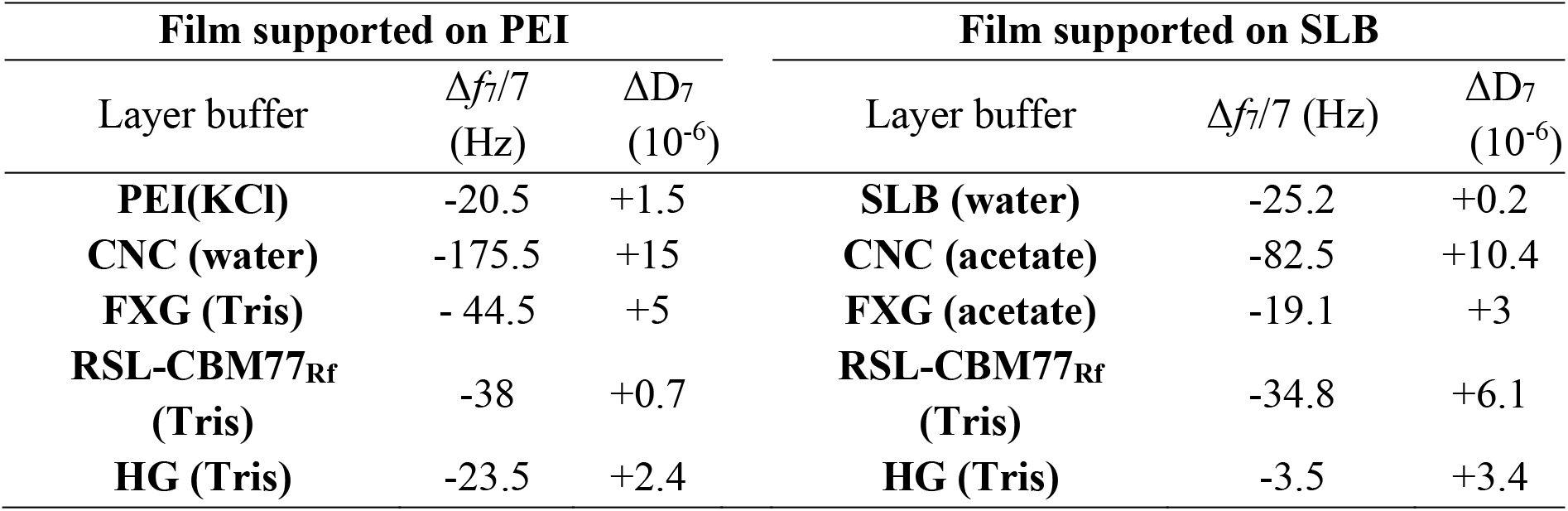
Frequency and dissipation variation quantified from experiments displayed in Figure 6.

The addition of solution of nanocrystals generates a large shift in frequency of -175.5 Hz indicating very strong interaction with the underlying polymer cushion, with slight changes upon rinsing with H_2_O and then Tris buffer. The further addition of FXG leads to an additional decrease of -44.5 Hz, again insensitive to rinsing. The important changes in the dissipation after the addition of both CNCs and FXG close to the aforementioned indicate that the layers are highly swollen with water, a characteristic that is expected from a loose interaction between XG and CNC, comparable to what is observed in the plant cell walls. When introducing the Janus lectin RSL-CBM77_Rf_, a third important decrease of the frequency (- 38 Hz) is observed. Interestingly, the level of this decrease is comparable to what is observed for FXG, meaning similar mass adsorption, but with lower dissipation, indicating strong interaction on this stable cushion, and stable after rinsing with the buffer. Finally, when HG was added, a fourth decrease of -23.5 Hz was observed with a slight increase of the dissipation. Indeed, HG is now interacting with the exposed CBM77_Rf_ as RSL did not exhibit any affinity with HG (see above).

In the second experiment, the multilayer film was built similarly but on a supported lipid bilayer (SLB) of DOPC that more closely mimics natural plant cell wall in comparison with the PEI. The SLB was formed by deposition of DOPC vesicles that open on the glass and form a lipid bilayer. The CNCs were then added as described previously resulting in a strong frequency change (-82.5 Hz). The amplitude was smaller than the one obtained on PEI but in agreement with a previous study on SLB as the net charge is higher in PEI layers than in SLB, inducing stronger interaction with CNCs (Y. Navon et al., 2020). Similarly, the deposition of FXG resulted in a clear frequency shift of -19.1 Hz that did not change in both cases after rinsing with acetate buffer, indicating a strong affinity. In this particular case, a necessary change of buffer (from acetate to Tris as protein is not active at low pH) creates rearrangement of the layers attested by an increase in the frequency and dissipation. However, the addition of RSL-CBM77_Rf_ did result in a frequency shift of -34.8 Hz equivalent to the one measured with the PEI-functionalized surface. Despite this nice proof of interaction, a smaller variation of -3.5 Hz is finally obtained when adding the homogalacturonan solution. At this stage, one has to conclude that the quality of the layers is a key factor when dealing with quantitative measurements, and if the PEI-supported proved to be a stable and reliable precursor, the SLB has to be optimized, for example, the nature of the lipids, in order to avoid the destabilization of the layers as observed in Figure 6B.

Control experiments were performed with RSL instead of Janus lectin, resulting in smaller step in frequency upon addition, consistent with its lower molecular weight. The absence of the CBM77_Rf_ moiety on the lectin resulted in total absence of binding of homogalacturonan for the last layer (Figure 7). The other control experiments made use of non fucosylated XG instead of FXG. In this case, no change in frequency was observed after RSL-CBM77_Rf_ injection confirming the need of fucose for the addition of this layer (Figure 7).

**Figure 7:**
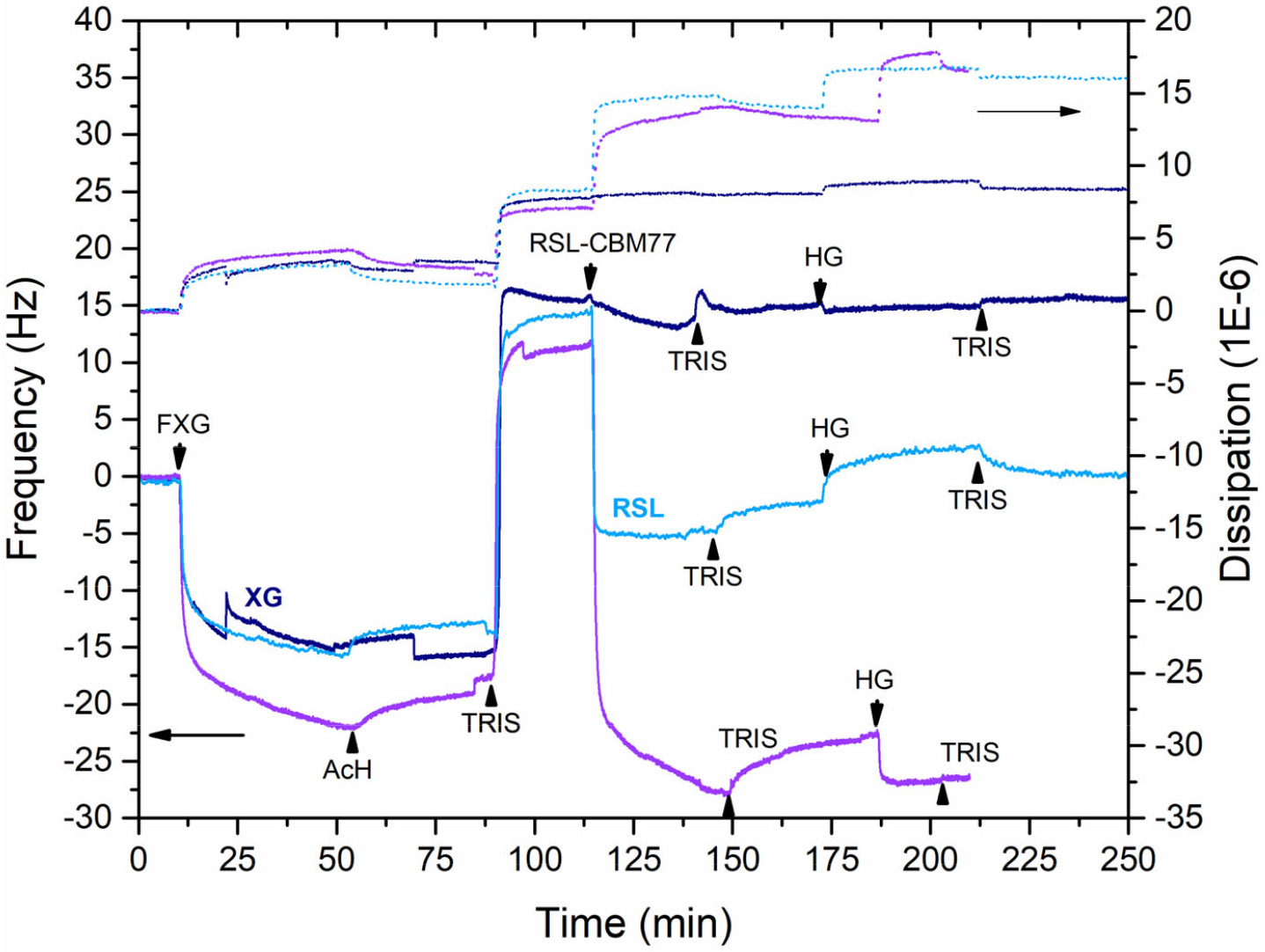
Overlay of control experiments. All the experiments were performed on multilayer constructure on SLB whereas either Janus lectin RSL-CBM77_Rf_ was replaced with RSL (light blue) and no binding of HG is observed in the last step or the nonfucosylated XG (dark blue) was used instead of FXG and in this case, no binding of RSL-CBM77_Rf_ nor of HG was observed. The original data with FXG and RSL-CBM77_Rf_ are shown in violet.

## Discussion

In the development of synthetic biology, the obtainment of artificial cells is a goal pursued by many teams. General efforts concentrate on the content of such cell, i.e. minimum genetic content to reproduce life, while it is also necessary to re-create surfaces of pseudo-cells with properties that are close enough to the real ones. Mimicking the surface of animal cells necessitates the incorporation of glycolipids, glycoproteins or other glycoconjugates to create the glycocalyx necessary for cell-cell interaction. In the case of plant cells, the complexity is much higher and polysaccharides with very different nature and properties must be assembled for obtaining the mechanical and biological properties obtained in Nature. We demonstrated here that the addition of a small fraction of well-designed protein participates in the building of cell wall mimics.

The diCBM77_Rf_ that has been constructed here has the ability to label strongly galacturonans, which could be of interest as a biomarker. Indeed, a limited number of antibodies are available as markers of polysaccharides in plants, and the molecular structure recognized by these antibodies often remains poorly characterized (Ruprecht et al., 2017). CBMs are more robust than antibodies but display weak affinity due to their monovalent character. Engineered diCBMs with specificities towards a component of plant cell walls may therefore represent useful tools for analyzing plant polysaccharides. Due to the large catalog of specificities toward plant polysaccharides exhibited by CBMs (Boraston et al., 2004; Duffieux et al., 2020), the concept can be extended towards other cell wall components.

This CBM77_Rf_ associated with lectin RSL resulted in a new tool for inducing the association of two soluble plant polysaccharides, i.e. pectin (homogalacturonan) and hemicellulose (fucosylated xyloglucan). The supported artificial plant cell wall that has been constructed is characterized by a single layer of cellulose nanocrystals on a lipid bilayer, with a thickness of 4-5 nm for SLB and 6-8 nm for CNCs in the case of a quasi-monolayer, according to previous work. FXG addition resulted in a more limited thickness of approximately 2 nm (Yotam Navon, 2020). The introduction of protein in the cell wall model allowed for completion with a thin pectin layer, but also influenced the flexibility, as indicated by analysis of the dissipation.

In conclusion, supported cell wall mimics that have been obtained in this study are of interest for research purposes. Cellulose-based containers built on lipid vesicles have already been described and completing such construct with different polysaccharides and interacting proteins is now possible. Indeed, solid support like cellulose films supported on polymer or lipid bilayers can be designed and were shown to exhibit high interaction level as in the solution state. Implementing biophysical experiments on heterogeneous substrates in the particular field of the plant cell wall is a step forward in the understanding of the interaction within such a complex system and open the routes to the design of new devices and innovative materials and a first step for the obtaining of artificial plant cell walls.

## Material and Methods

### Oligosaccharides and polymers preparation

Oligosaccharide XFGol was purchased from Elicityl (PEL101GV) and the reported 85% purity was checked by NMR analysis. GalA_DP7 was obtained from the CERMAV collection (Grenoble, France) (Gouvion et al., 1994). A purity check was performed by gel permeation chromatography indicating that the sample consists mainly of DP7 with limited contamination by DP8. 2-fucosyl lactose was obtained from CERMAV collection and the purity was checked by NMR analysis. Homogalacturonan (HG) from orange was obtained from Sigma (P3889) and further purified by dialysis in order to remove impurities and small polymers. The absence of the methyl ester functional group was verified by NMR analysis. Gel permeation chromatography on SB 806 M HQ column indicated an MW of 35 kDa. Nonfucosylated xyloglucan from tamarin fruit (Glyloid 3S, Dainippon) was further purified by ethanol in order to remove the contaminants. Fucosylated xyloglucan (FXG) was obtained from apple pomace, inspired by (Watt et al., 1999). Gel permeation chromatography on SB 806 M HQ column gave a molecular weight of 55 kDa for FXG and acidic hydrolysis treatment followed by anion-exchange liquid chromatography indicated that the fucose content is 5% (Table 4).

**Table 4:** Monosaccharide composition of apple FXG presented as mass percentage.

|  | **Fuc** | **Rha** | **Ara** | **Gal** | **Glc** | **Man** | **Xyl** | **Total** |
| --- | --- | --- | --- | --- | --- | --- | --- | --- |
| **%** | 5.30 | 3.71 | 3.85 | 10.85 | 29.49 | 9.05 | 21.39 | 83.63 |
| **mass\*** | ± 0.32 | ± 0.13 | ± 0.10 | ± 0.61 | ± 0.94 | ± 0.23 | ± 0.80 | ± 3.14 |

\* the mass percentage corresponds to monosaccharide mass (in g) in 100 g of sample.

The CNCs suspensions were prepared by acid hydrolysis of cotton linters (Buckeye Cellulose Corporation) according to reported method (Dong et al., 2998) in the form of a suspension at 2.47% wt obtained by hydrolysis of cotton fiber with sulfuric acid. This treatment has the particularity to graft sulfate groups on the CNC, covering them with negative charges. The CNC suspension was subsequently sonicated for 30 s at 10% and then filtered using a 0.45 µm syringe filter.

### Gene design and cloning

The original amino acid sequence of the carbohydrate-binding module CBM77_Rf_ from *Ruminococcus flavefaciens* was obtained from the PDB database. The gene *dicbm77*_*Rf*_ was designed as a tandem repeat of two individual CBM77_Rf_ domains connected *via* ALNGSELGSGSGLSSLGEYKDI linker. The gene *rsl-cbm77*_*Rf*_ was designed as a fusion chimera with RSL at N-terminus and CBM77_Rf_ at C-terminus *via* linker GGGGSGGGGS. The genes were ordered from Eurofins Genomics (Ebersberg, Germany) after codon optimization for the expression in the bacteria *Escherichia coli*. The restriction enzyme sites of NdeI and XhoI were added at 5’ and 3’ ends, respectively. The synthesized genes were delivered in plasmid pEX-A128-diCBM77_Rf_ and pEX-A128-RSL-CBM77_Rf_ respectively. The plasmid pEX-A128-diCBM77_Rf_ and the pET-TEV vector (Houben et al., 2007) were digested with the NdeI and XhoI restriction enzymes to ligate *dicbm77*_*Rf*_ in pET-TEV to fuse a 6-His Tag cleavable with TEV protease at the *N*-terminus of diCBM77_Rf_. Similarly, plasmids pEX-A128-RSL-CBM77_Rf_ and the pET-25b+ were digested with the NdeI and XhoI restriction enzymes to ligate *rsl-cbm77*_*Rf*_ into pET-25b+. After transformation by heat shock in *E. coli* DH5α strain, a colony screening was performed, and the positive plasmids were amplified and controlled by sequencing.

### Protein expression

*E. coli* BL21(DE3) cells were transformed by heat shock with pET-TEV-diCBM77_Rf_ plasmid prior pre-culture in Luria Broth (LB) media with 25 μg/mL kanamycin at 37°C under agitation at 180 rpm overnight. The next day, 10 mL of pre-culture was used to inoculate 1 L LB medium with 25 μg/mL kanamycin at 37°C and agitation at 180 rpm. When the culture reached OD_600nm_ of 0.6 - 0.8, the protein expression was induced by adding 0.1 mM isopropyl β-D-thiogalactoside (IPTG), and the cells were cultured at 16°C for 20 hours.

Correspondingly, *E. coli* KRX cells were transformed by heat shock with pET-25b+-RSL-CBM77_Rf_ plasmid and pre-cultured in LB media substituted with 50 μg/mL ampicillin at 37°C under agitation at 180 rpm overnight. The following day, 10 mL of pre-culture was used to inoculate 1 L LB medium with 50 μg/mL ampicillin at 37°C and agitation at 180 rpm. When reached OD_600nm_ of 0.6 - 0.8, the protein expression was induced by adding 1% L-rhamnose, and the cells were cultured at 16°C for 20 hours.

The cells were harvested by centrifugation at 14000 × *g* for 20 min at 4°C and the cell paste was resuspended in 20 mM Tris/HCl pH 7.5, 100 mM NaCl (Buffer A), and lysed by a pressure cell disruptor (Constant Cell Disruption System) with a pressure of 1.9 kBar. The lysate was centrifuged at 24 000 × *g* for 30 min at 4°C and filtered on a 0.45 µm syringe filter prior to loading on an affinity column.

### Protein purification

#### diCBM77_Rf_

The cell lysate was loaded on 1 mL HisTrap column (Cytiva) pre-equilibrated with Buffer A. The column was washed with Buffer A to remove all contaminants and unbound proteins. The diCBM77_Rf_ was eluted by Buffer A in steps during which the concentration of imidazole was increased from 50 mM to 500 mM. The fractions were analyzed by 12% SDS PAGE and those containing diCBM77_Rf_ were collected and deprived of imidazole by dialysis in Buffer A. N-terminal His-tag was removed by TEV cleavage with the ratio 1:50 mg of TEV:protein in the presence of 0.5 mM EDTA and 1mM TCEP over night at 19°C. After, the protein mixture was purified on 1 mL HisTrap column (Cytiva) and the pure protein was concentrated by Pall centrifugal device with MWCO 10 kDa and stored at 4°C.

#### RSL-CBM77_Rf_

Likewise, the cell lysate was loaded on 10 mL D-mannose-agarose resin (Merck) pre-equilibrated with Buffer A. The column was washed with Buffer A to remove all contaminants and unbound proteins and the flow-through was collected. RSL-CBM77_Rf_ was eluted by Buffer A with the addition of 100 mM D-mannose in one step. Due to insufficient binding capacity of the column, the flow-through was reloaded on the column several times and the protein was eluted as described previously. The fractions were analyzed by 12% SDS PAGE and those containing RSL-CBM77_Rf_ were collected and dialyzed against Buffer A. The protein was concentrated by Pall centrifugal device with MWCO 10 kDa and the pure protein fractions were pooled, concentrated, and stored at 4°C.

### Isothermal Titration Calorimetry (ITC)

ITC experiments were performed with MicroCaliTC200 (Malvern Panalytical). Experiments were carried out at 25°C ± 0.1°C. Proteins and ligands samples were prepared in Buffer A, except of oligosaccharide GalA_DP7 which was prepared in 20 mM Tris pH 7.5. The ITC cell contained proteins in a concentration range from 0.05 mM to 0.2 mM. The syringe contained the ligand solutions in a concentration from 50 μM to 10 mM. 2 μL of ligands solutions were injected into the sample cell at intervals of 120 s while stirring at 750 rpm. Integrated heat effects were analyzed by nonlinear regression using one site binding model (MicroCal PEAQ-ITC Analysis software). The experimental data were fitted to a theoretical curve, which gave the dissociation constant (Kd) and the enthalpy of binding (∆H).

### Decomposition analysis of FXG

Apple FXG 4 mg was placed in a dry bath in the presence of 1 mL of 2N TFA, for 4 h at 110°C. After cooling, the sample was filtered on 0.2 μm then diluted 40 times in reverse osmosis water. Subsequently, the sample was injected on anion exchange liquid chromatography, on a DIONEX ICS6000 system equipped with a CarboPac PA1 pre-column and column (250*2 mm) and a PAD (pulsed amperometry) detector. The injected sample volume was 5 μL and the purification run at 28°C. The elution was carried out at 0.25 mL/min under the following conditions: T0 to T18 min: NaOH 0.016 M, T18 to T20 min: NaOH 0.016 M + linear gradient in NaOAc from 0 to 0.02 M, T20 to T25: linear gradient in NaOH from 0.016 M to 0.2 M and linear gradient in NaOAc from 0.02 M to 0.5 M, T25.1 to T30: NaOH 0.1 M + NaOAc 0.5 M. A calibration curve for each monosaccharide (fucose, rhamnose, arabinose, galactose, glucose, mannose, xylose, ribose, galacturonic acid, glucuronic acid was performed (2*10^−3^ to 3*10^−2^ mg/mL) in the same operating conditions. The sample was injected three times.

### Quartz crystal microbalance with dissipation monitoring (QCM-D) measurement

QCM-D measurements were performed using Q-Sense E4 instruments (Biolin Scientific) equipped with one to four flow modules. Silica-coated AT-cut quartz crystals (QSX 303) were purchased from Biolin Scientific (Stockholm, Sweden). Besides measurement of bound mass, which is provided from changes in the resonance frequency, *f*, of the sensor crystal, the QCM-D technique also provides structural information of biomolecular films via changes in the energy dissipation, *D*, of the sensor crystal. *f* and *D* were measured at the fundamental resonance frequency (4.95 MHz) as well as at the third, fifth, seventh, ninth, eleventh, and thirteenth overtones (*i* = 3, 5, 7, 9, 11 and 13). Normalized frequency shifts D*f* = D*f*_*i*_/*i* and dissipation shifts Δ*D* = Δ*D*_*i*_ corresponding to the seventh overtone only were presented for clarity.

The sensors were rinsed with a 2 wt % SDS aqueous solution followed by abundant water rinsing and dried under a nitrogen stream. The sensors were then exposed to a UV/ozone lamp for 10 min prior to the measurements.

Flow rate was controlled using a peristaltic pump (ISM935C, Ismatec, Switzerland). The temperature of the E4 QCM-D platform and all solutions were stabilized to ensure stable operation at 24 °C. All buffers were previously degassed in order to avoid air bubble formation in the fluidic system. All experiments were performed in duplicates.

Polyethylene-imine (PEI), a branched cationic polymer which can serve as a good primer was injected (1 g. L^-1^) in water. In all the experiments done with PEI, the tubing diameter was 0.25 mm and the flow rate was set at 10 µL.min^-1^. Second injection was performed with CNC (1 g. L^-1^) then FXG (1 g. L^-1^) in pure water. This was followed by rinsing with 20 mM Tris (pH=7.5), 100 mM NaCl and injection of RSL-CBM77_Rf_ (different concentrations up to 10 µM) and then HG (1 g. L^-1^) in the same buffer. Nonfucosylated XG from tamarin or RSL were used in control experiments.

To produce an SLB on QCM-D, a small unilamellar vesicles (SUVs) suspension was injected into the measurement chamber. The DOPC (1,2-dioleoyl-sn-glycero-3-phosphocholine) SUVs were prepared as previously described (Y. Navon et al., 2020) and injected in 100 mM acetate (pH=3). With SLB surface was done using tubing with a 0.75 mm diameter and a flow of 50 µL.min^-1^. CNC (1 g. L^-1^) was then injected in the same buffer followed by FXG (1 g. L^-1^) then rinsed with 20 mM Tris (pH=7.5). This buffer was also used for injection of RSL-CBM77_Rf_ (10 µM) and HG (1 g. L^-1^).

## Acknowledgements

This research was funded by the European Union Horizon 2020 Research and Innovation Program under the Marie Skłodowska-Curie grant agreement synBIOcarb (No. 814029). The authors thank the nano-bio platform at ICMG, DCM, University of Grenoble Alpes (ICMG FR 2607) for the use of QCM-D as well as the Glyco@Alps program (Investissements d’Avenir Grant No. ANR-15-IDEX-02) and Labex Arcane/CBH-EUR-GS (ANR-17-EURE-0003). The authors acknowledge PCANS platform at Cermav CNRS, especially Laurine Buon and Eric Bayma for the oligo and polysaccharide characterization. We would also like to thank Microbiology platform at Cermav CNRS for providing 2-fucosyl lactose.

